## Supplemental Figures for "α-Synuclein strains that cause distinct pathologies differentially inhibit proteasome": SupFig.pdf

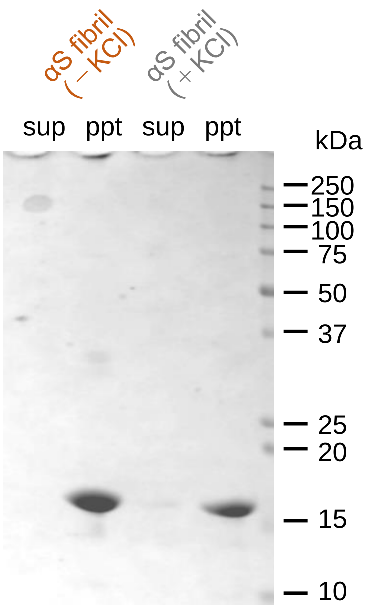


**Suzuki et al., Figure 1-figure supplement 1**

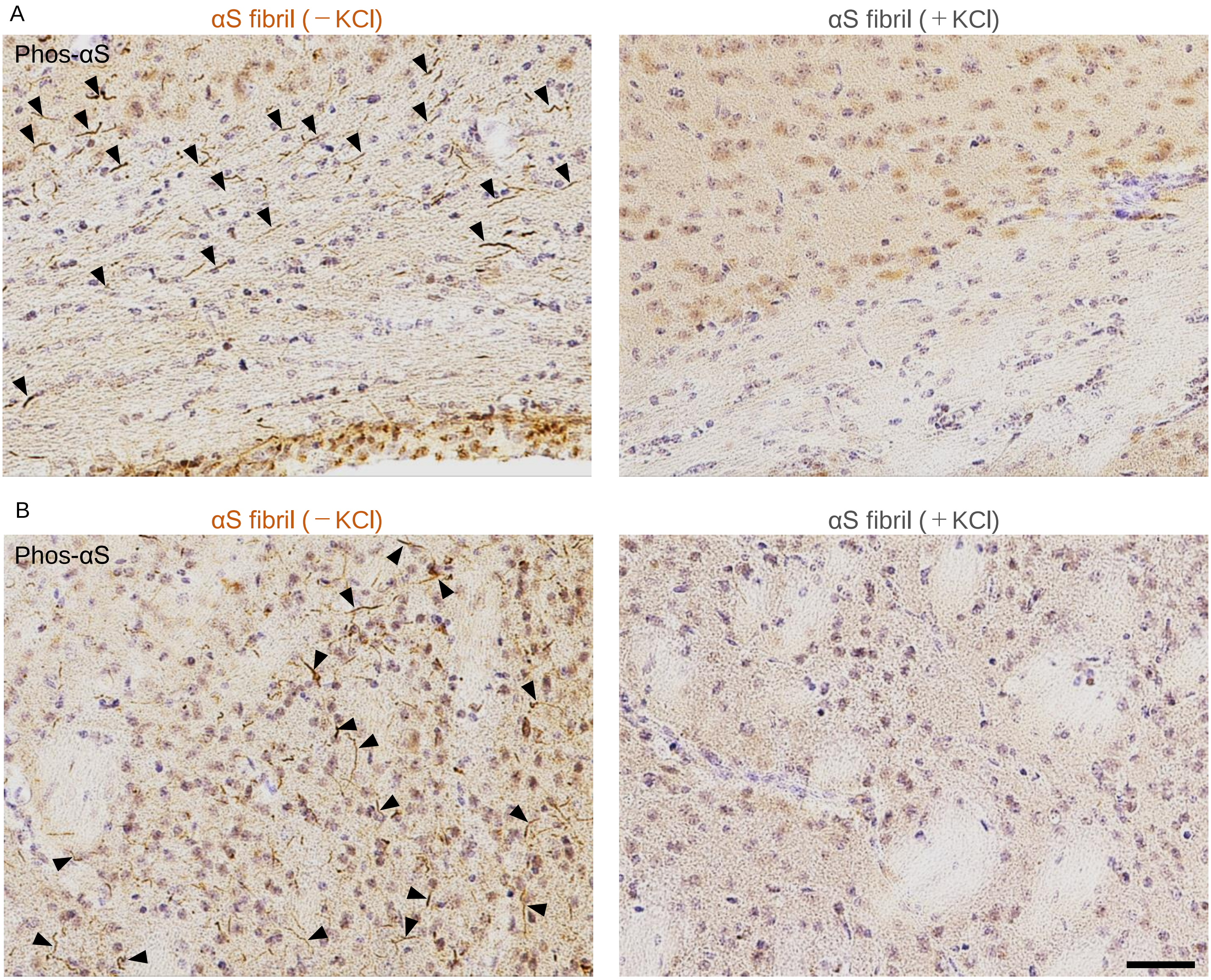


**Suzuki et al., Figure 2-figure supplement 1**

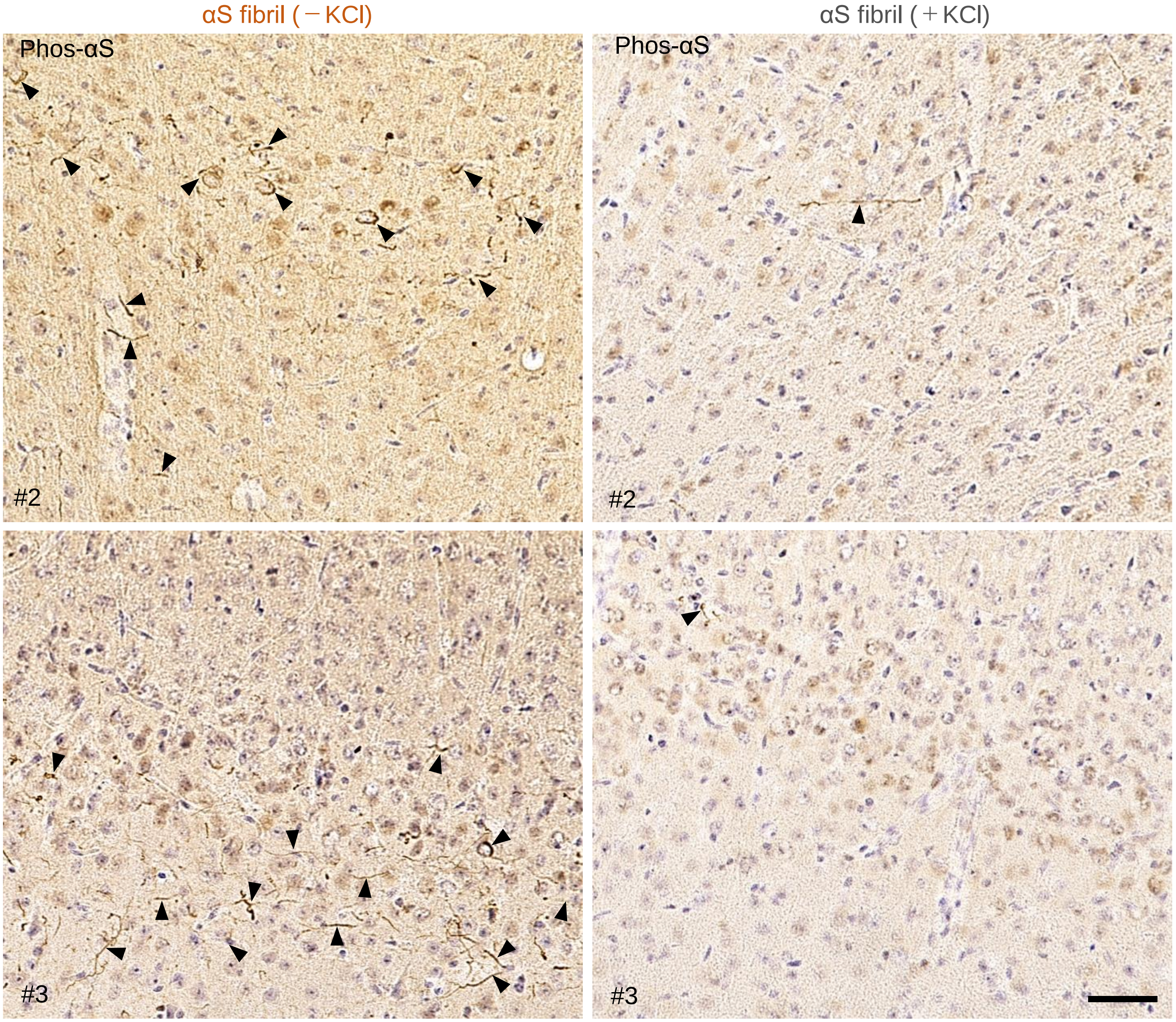


**Suzuki et al., Figure 2-figure supplement 2**

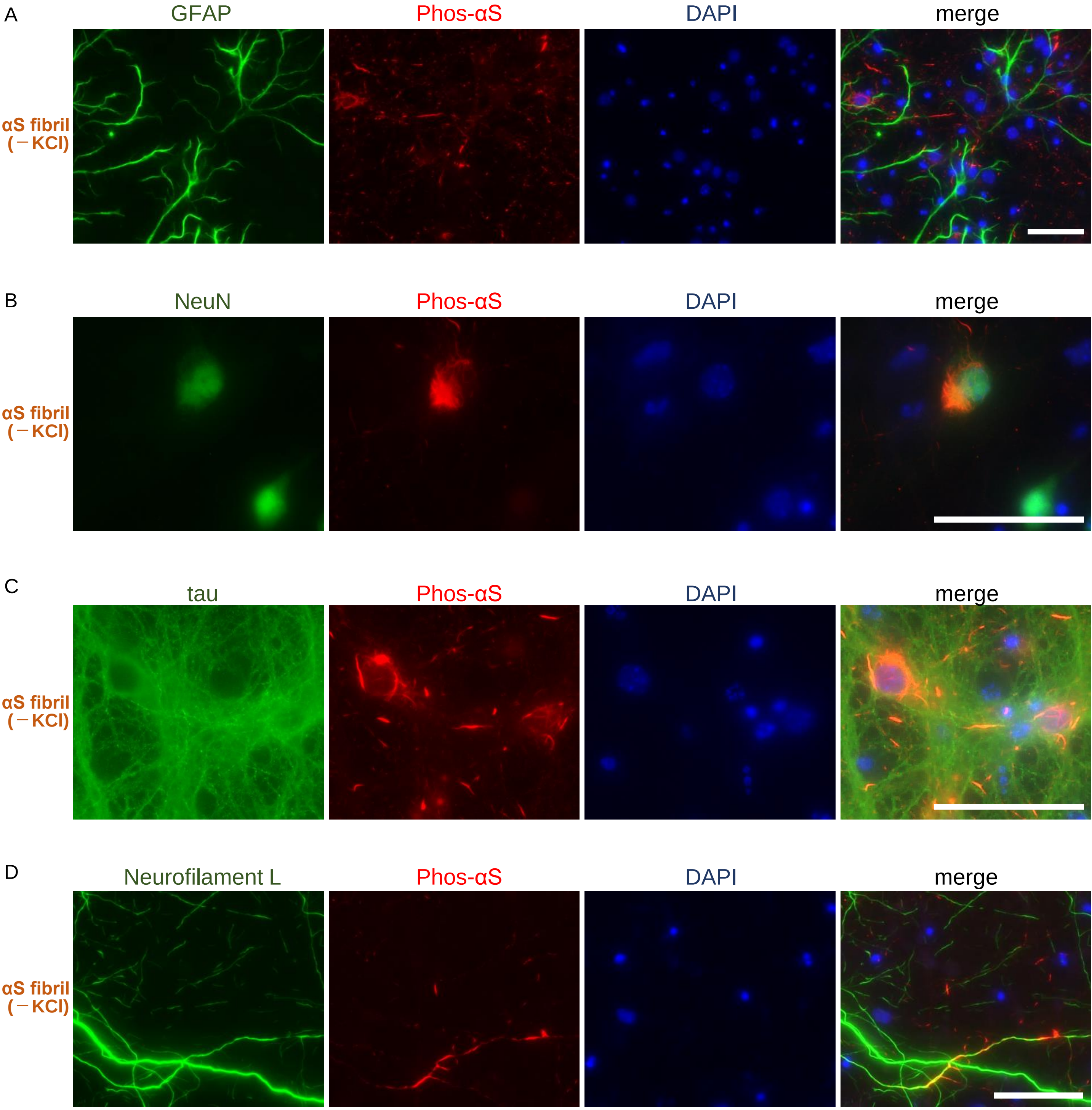


**Suzuki et al., Figure 3- figure supplement 1**

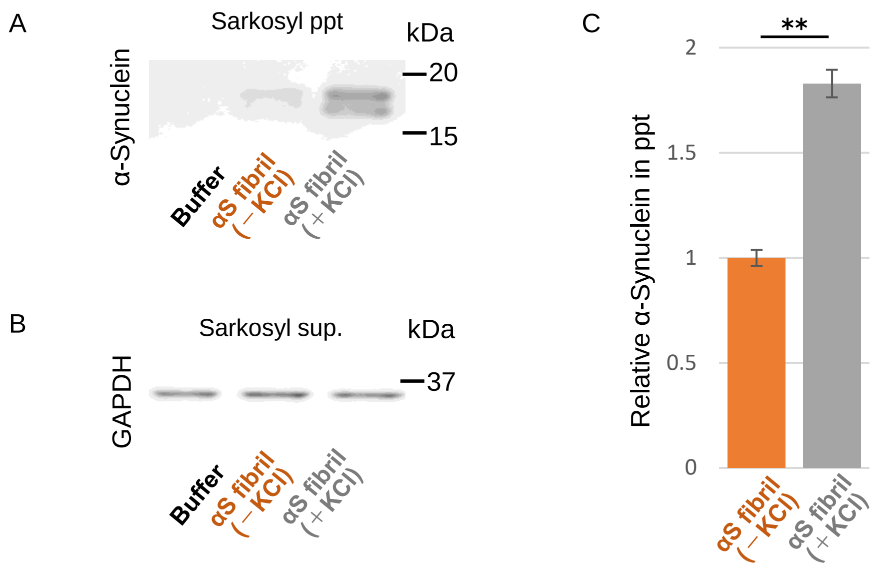


**Suzuki et al., Figure 3- figure supplement 2**

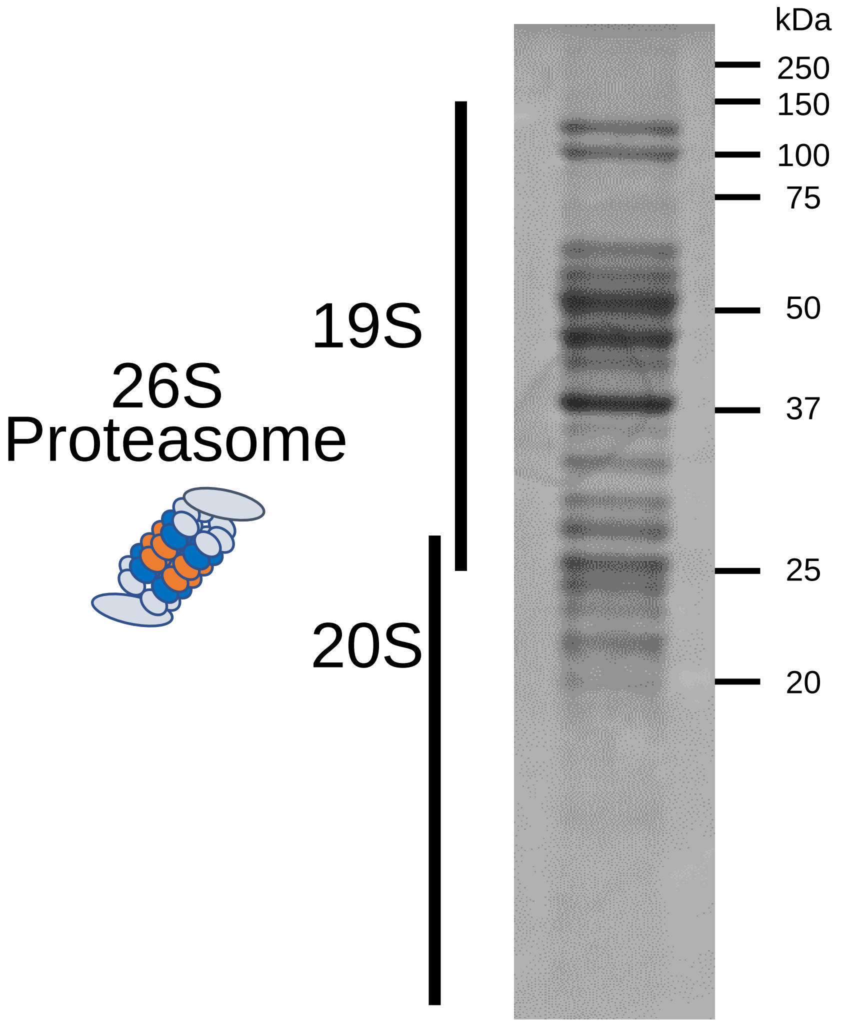


**Suzuki et al., Figure 4-figure supplement 1**
